# Collateral sensitivity and cross resistance in six species of bacteria exposed to six classes of antibiotics

**DOI:** 10.1101/2024.12.12.628210

**Authors:** Xinyu Wang, Luyuan Nong, Gioia Schaar, Martijs Jonker, Wim de leeuw, Benno H. ter Kuile

**Affiliations:** Biology and Microbial Food Safety, Swammerdam Institute for Life Sciences, University of Amsterdam, Amsterdam, The Netherlands; RNA Biology & Applied Bioinformatics, Swammerdam Institute for Life Sciences, University of Amsterdam, Amsterdam, The Netherlands

**Keywords:** *fusA* mutation, collateral sensitivity, cross resistance, convergent evolution, kanamycin resistance

## Abstract

*De novo* resistance can be developed in bacteria because of exposure to sublethal concentrations of antibiotics. Once the strain has become resistant to an initial antibiotic, this can cause cross-resistance or collateral sensitivity to a second antimicrobial. Specific collateral sensitivity is rarely conserved across species because the mechanisms triggered in different microorganisms to resist antibiotics are often different. In this study, we explored which collateral sensitivity or cross resistance networks are present in six species of bacteria with induced *de novo*-resistance. These six species were induced to become resistant to amoxicillin/cefepime, enrofloxacin, kanamycin, tetracycline, erythromycin and chloramphenicol(1). After that, the collateral sensitivity and the cross-resistance networks were evaluated by measuring increase or decrease of MIC of thirteen antibiotics that are often used in the clinic. Collateral sensitivity for kanamycin occurred in five species of strains resistant to chloramphenicol and tetracycline and for β-lactam in strains of five species resistant to kanamycin. Further genetic analysis clarified that *fusA* consistently mutated in five species of bacteria made *de novo* resistant against kanamycin, suggesting that *fusA* operates in parallel with the other mechanisms related to antimicrobial resistance Based on considerations of resistance, a treatment protocol starting with chloramphenicol/tetracycline, followed by kanamycin and ending with amoxicillin may eliminate bacteria that have developed resistance against the initial treatment.

## Introduction

Antibiotic-resistant bacterial infections have become a serious public health concern across the world(2). When bacteria initially become resistant to one antibiotic, this may increase or decrease the susceptibility to other antibiotics compared to naïve wild type strain, defined as collateral sensitivity (CS) and cross-resistance (CR), respectively(3). Cross-resistance can occur due to multi-target mechanisms, such as efflux pumps(4–6). Persistence of CR hampers clearing pathogens that have become resistant to the affected antimicrobials. Inversely, collateral sensitivity might be used to advance treatment options. In that case, alternative treatments could be developed based on CS to eliminate bacteria resistant to one antibiotic using specifically designed treatments with other compounds(7, 8). Antibiotic combinations that take advantage of CS could be designed to be more effective than the individual antibiotics alone, thereby reducing development of resistance(7, 9).

Both CS and CR have been described in a variety of bacterial species and using CS is regarded as a viable technique to combat antibiotic resistance(10). Resistance to ciprofloxacin, a fluoroquinolone, increased susceptibility to the aminoglycoside tobramycin and to the β-lactam aztreonam in *Pseudomonas aeruginosa*(7, 11). Similarly, tobramycin resistance increased the susceptibility for phosphomimic acid but not for ciprofloxacin(12). Collateral sensitivity networks have been identified in *Escherichia coli*, where aminoglycoside resistance was linked to the AcrAB efflux system(13). The diverse genomic backgrounds enable bacteria to apply mutational trajectories to resist a specific antibiotic, leading to differences in phenotype, causing CS. The most notable case is the activation of efflux pumps in gram negative of bacteria, but this is less common in gram positives. The CS in multiple species has the potential to increase the effectiveness of antibiotic treatments in hospitals(10).

This study aimed at identifying the CS and CR networks in six species of bacteria made resistant against in total thirteen antibiotics. *Staphylococcus aureus, Enterococcus faecalis, Bacillus subtilis, Yersinia enterocolitica, Salmonella enterica* and *Acinetobacter pittii* were induced to evolve *de novo* resistance against six different classes of antibiotics: Beta-lactam, fluoroquinolones, aminoglycosides, tetracyclines, macrolides and chloramphenicols(1). No species was made resistant against more than six antibiotics, avoiding compounds against which a species was inherently resistant.

The following questions were addressed: To which extent do collateral sensitivity and cross-resistance occur after development of *de novo* resistance? Can patterns of CS be observed across species? Are the appearance of CS and CR consistent enough to function as a foundation for development of strategies to prevent antimicrobial resistance?

## Results

In order to identify patterns in the occurrence of collateral sensitivity (CS) and cross resistance (CR), six species of bacteria were made resistant against six antibiotics and tested against in total 13 antibiotics. Because of inherent resistance, not all combinations of drug and species were relevant (Figure 1). CS and CR are visualized for six bacterial species made *de novo* resistant against at least one of the six classes mentioned in the introduction. At first view, cross-resistance seems to occur more often than collateral sensitivity. This conclusion is incorrect, because in these heatmaps the original antibiotic against which resistance was developed, is also included, sometimes accompanied by chemically closely related compounds. Collateral sensitivity between chemically unrelated antibiotics does occur more often than CR (Figure 1). Strong CR occurred between fluoroquinolone resistance and induced cefepime resistance in *S. aureus* and *E. faecalis*; The cefepime and amoxicillin induce more often CS than the other antimicrobials tested. The aminoglycosides used in this study show mild CS with several other antimicrobials across species. The strains of *S. enterica* exposed to amoxicillin, *A. pittii* to enrofloxacin, *E. faecalis* to kanamycin, *S. aureus* to tetracycline, *S. enterica* to erythromycin and *S. aureus* to chloramphenicol all barely developed resistance. Consequently, they did not develop noticeable CS and CR.

**Figure 1.**
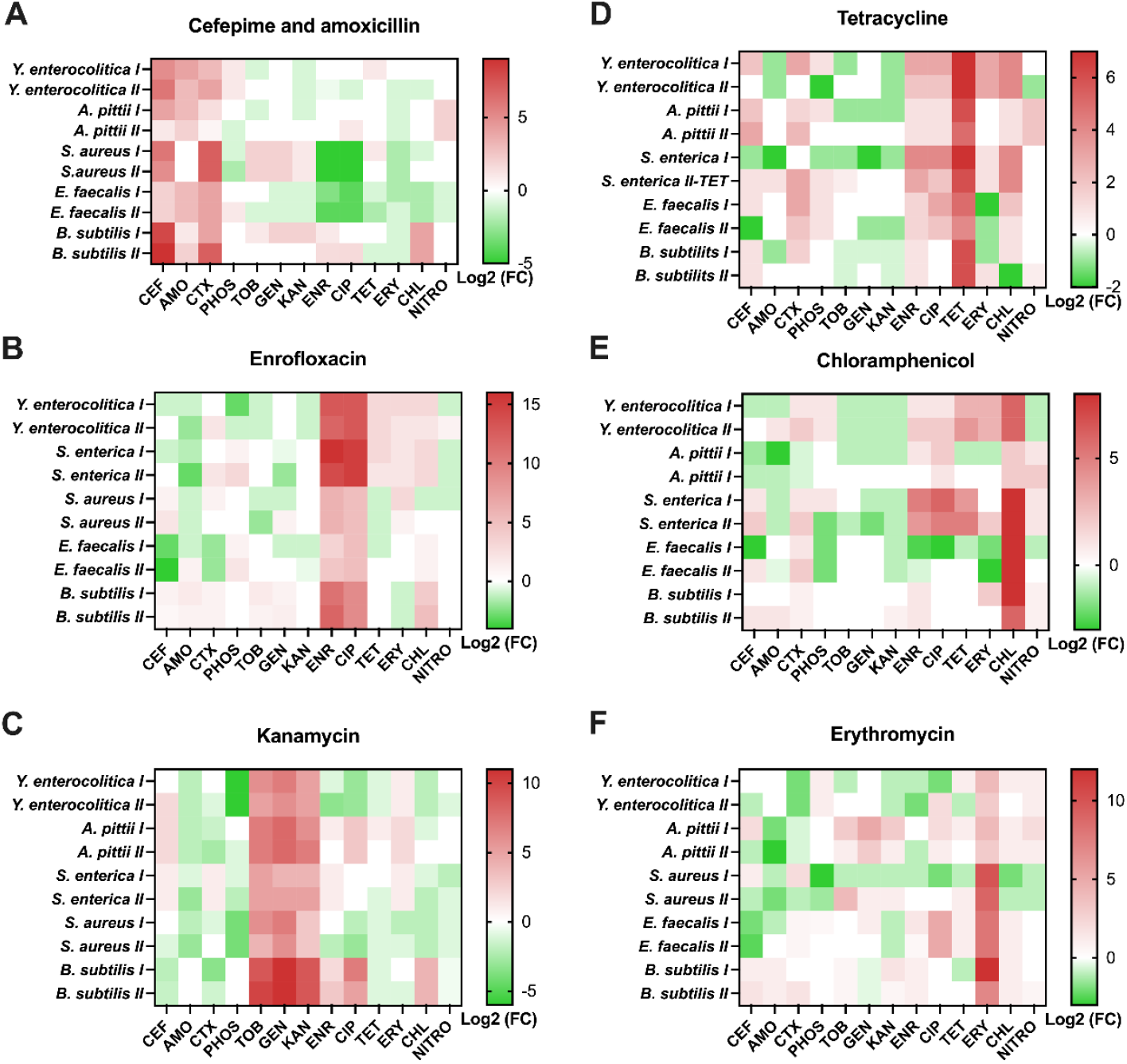
Visualization of collateral sensitivity and cross-resistance and cross-resistance of strains made resistant to **A**) amoxicillin/cefepime, **B**) enrofloxacin, **C**) kanamycin, **D**) tetracycline **E**) chloramphenicol **F**) erythromycin. The intensity of red represents the degree of cross-resistance and the intensity of green the degree of collateral sensitivity based on the Log2 of the foldchange in MIC-values of Table 2.

One of the most compelling findings is the conserved CS observed in five strains resistant to tetracycline/chloramphenicol, kanamycin, and amoxicillin/cefepime. Five species of bacteria resistant to tetracycline/chloramphenicol did not show any CR to aminoglycosides. Instead, they consistently exhibited slight or clear CS to aminoglycoside antibiotics. Additionally, the strains resistant to kanamycin displayed CR to gentamicin and tobramycin while consistently maintaining CS to both amoxicillin and cefepime, revealing a reliable solution to clear out the resistant strain in these five species (Figure 1).

The convergence of kanamycin resistance across five species can primarily be attributed to a crucial common mutation in the *fusA* gene, which drives most kanamycin resistance(1). The fusA gene codes for the elongation factor G in E. coli, which is important for bacterial protein synthesis and is associated with drug resistance in Clostridium difficile(14). This gene was mutated in all replicates during development of kanamycin resistance (table 1). The number of times that an alanine positioned in the 500’s was replaced is striking (7 times). Most of times the new amino acid was valine. The reverse only happened once, at a very different location (86). Around the location of alanine replacement, glycine (5 X) and phenylalanine (6 X) were almost as often replaced. Glycine most often by cysteine and phenylalanine by leucine, isoleucine or valine. Otherwise, only 2 serine residues are replaced, a single threonine and one valine. All the replacements are very logical if the codon’s are taken into consideration and all can be the result of a single nucleotide replacement. However, the overall effect on kanamycin resistance is still striking, so the selective advantages must have been considerable.

**Table 1.**
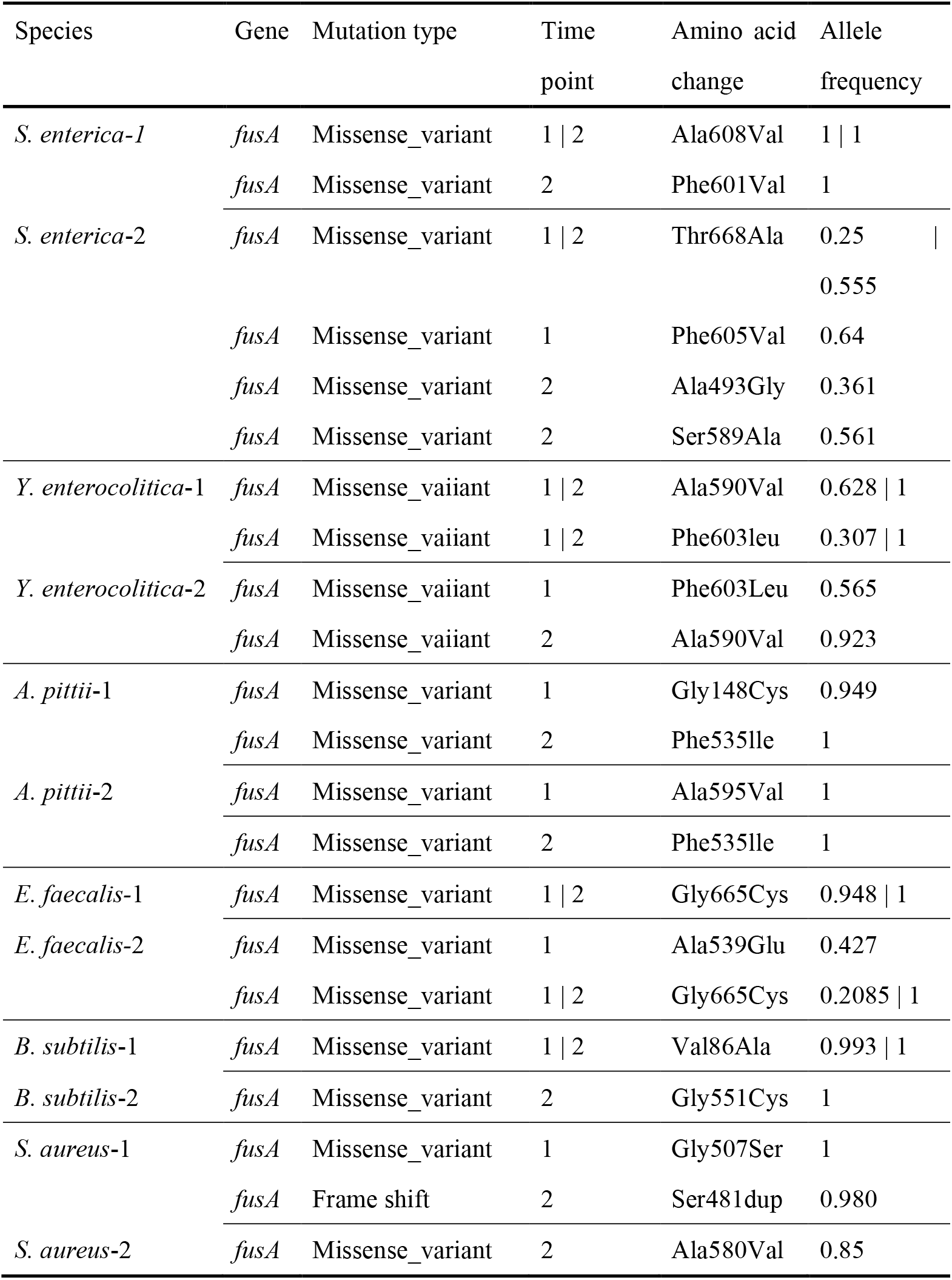
Mutations of *fusA* in six species as a result of induced kanamycin resistance.

| Species | Gene | Mutation type | Time point | Amino acid change | Allele frequency |
| --- | --- | --- | --- | --- | --- |
| *S. enterica*-1 | *fusA* | Missense_variant | 1 \| 2 | Ala608Val | 1 \| 1 |
|  | *fusA* | Missense_variant | 2 | Phe601Val | 1 |
| *S. enterica*-2 | *fusA* | Missense_variant | 1 \| 2 | Thr668Ala | 0.25 \| 0.555 |
|  | *fusA* | Missense_variant | 1 | Phe605Val | 0.64 |
|  | *fusA* | Missense_variant | 2 | Ala493Gly | 0.361 |
|  | *fusA* | Missense_variant | 2 | Ser589Ala | 0.561 |
| *Y. enterocolitica*-1 | *fusA* | Missense_vaiiant | 1 \| 2 | Ala590Val | 0.628 \| 1 |
|  | *fusA* | Missense_vaiiant | 1 \| 2 | Phe603leu | 0.307 \| 1 |
| *Y. enterocolitica*-2 | *fusA* | Missense_vaiiant | 1 | Phe603Leu | 0.565 |
|  | *fusA* | Missense_vaiiant | 2 | Ala590Val | 0.923 |
| *A. pittii*-1 | *fusA* | Missense_variant | 1 | Gly148Cys | 0.949 |
|  | *fusA* | Missense_variant | 2 | Phe535Ile | 1 |
| *A. pittii*-2 | *fusA* | Missense_variant | 1 | Ala595Val | 1 |
|  | *fusA* | Missense_variant | 2 | Phe535Ile | 1 |
| *E. faecalis*-1 | *fusA* | Missense_variant | 1 \| 2 | Gly665Cys | 0.948 \| 1 |
| *E. faecalis*-2 | *fusA* | Missense_variant | 1 | Ala539Glu | 0.427 |
|  | *fusA* | Missense_variant | 1 \| 2 | Gly665Cys | 0.2085 \| 1 |
| *B. subtilis*-1 | *fusA* | Missense_variant | 1 \| 2 | Val86Ala | 0.993 \| 1 |
| *B. subtilis*-2 | *fusA* | Missense_variant | 2 | Gly551Cys | 1 |
| *S. aureus*-1 | *fusA* | Missense_variant | 1 | Gly507Ser | 1 |
|  | *fusA* | Frame shift | 2 | Ser481dup | 0.980 |
| *S. aureus*-2 | *fusA* | Missense_variant | 2 | Ala580Val | 0.85 |

**Table 2.**
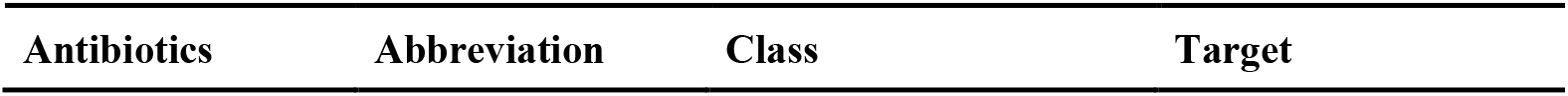
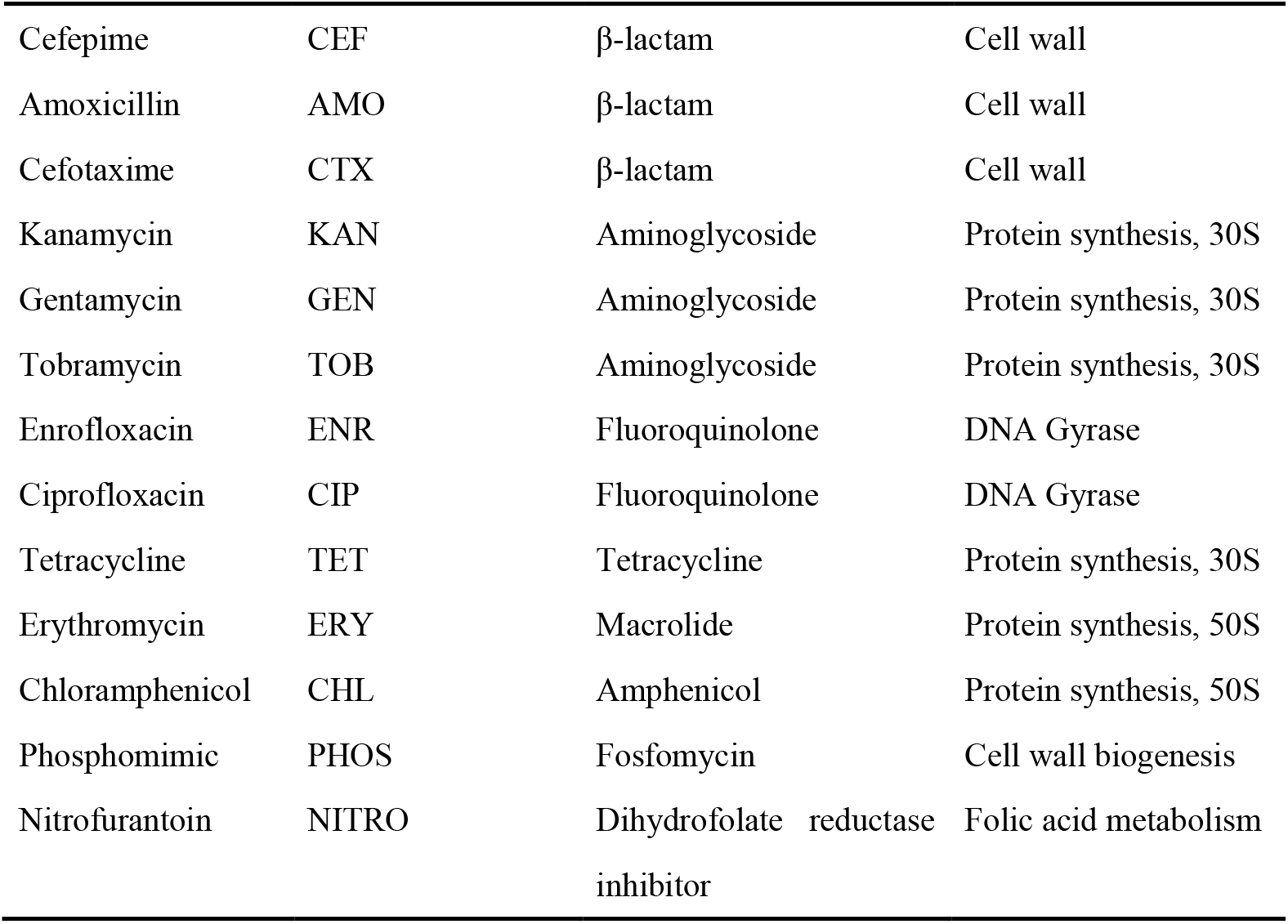
List of antibiotics used in the collateral sensitivity testing.

| Antibiotics | Abbreviation | Class | Target |
| --- | --- | --- | --- |
| Cefepime | CEF | β-lactam | Cell wall |
| Amoxicillin | AMO | β-lactam | Cell wall |
| Cefotaxime | CTX | β-lactam | Cell wall |
| Kanamycin | KAN | Aminoglycoside | Protein synthesis, 30S |
| Gentamycin | GEN | Aminoglycoside | Protein synthesis, 30S |
| Tobramycin | TOB | Aminoglycoside | Protein synthesis, 30S |
| Enrofloxacin | ENR | Fluoroquinolone | DNA Gyrase |
| Ciprofloxacin | CIP | Fluoroquinolone | DNA Gyrase |
| Tetracycline | TET | Tetracycline | Protein synthesis, 30S |
| Erythromycin | ERY | Macrolide | Protein synthesis, 50S |
| Chloramphenicol | CHL | Amphenicol | Protein synthesis, 50S |
| Phosphomimic | PHOS | Fosfomycin | Cell wall biogenesis |
| Nitrofurantoin | NITRO | Dihydrofolate reductase inhibitor | Folic acid metabolism |

## Discussion

The molecular mechanisms causing collateral sensitivity are poorly understood(15). The effects of CS have primarily been investigated in the framework of resistance evolution of single laboratory species(16–18). The investigation of the convergence of CS phenotypes in six de novo resistant strains against thirteen antibiotics in this study focused on the mechanisms underlying CS. One can assume that the cellular adjustments and/or DNA mutations that result in resistance to one antibiotic make the cell more vulnerable to a second antimicrobial with a different mode of action. In this study CS occurred for various combinations of bacterial species and two antibiotics, the initial that induced resistance and the second that either showed CS or not (drugs/bug). Even when CS is reproducible for specific drugs/bug combinations, it is usually not conserved after growth for some time in the absence of the initial antibiotic evolution(19–21). This suggests that both CS and CR can be caused by adaptation at the level of cellular mechanisms, rather than DNA mutations. The trade-off between functions that cells make, such as maintaining pH or salt balance against pumping out an antibiotic, can cause either CR or CS. If an induced efflux pump is effective against two antibiotics with different modes of action, the result will be cross-resistance(22). If the efflux pump is not effective against the second drug, but still uses energy that is then not available for combatting the second antibiotic, CS will be the outcome.

Questions remain regarding the convergence of CS phenotypes across species. Recent studies examined single strains with various knockout backgrounds(12, 23, 24), clinical isolates from a single species(10, 17), and different *ESKAPE* pathogens(10, 25). Since to the evolutionary trajectory is limited by the genomic context, the CS phenotypes are rarely consistent between various species, leading treatment failure in the clinic(12, 24). However, the target mutations in the node of the evolution trajectory, which have a positive epistasis with the other mutations, are rarely absent in the set of acquired mutations, thereby shaping a robust resistance phenotype(7, 26). A discrete CS phenotype is observed across six species of bacterial investigation. CR is only conserved within strains resistant to the same class of antibiotics. In the preceding investigation(1), the unique evolutionary trajectory is mostly observed in six species of bacteria exposed to six antibiotics. Six strains developed high levels of antibiotic resistance at the end of evolution, but the genetic changes were mainly acquired from the diverse mechanisms. The genetic changes in resistant strains indicating the resistance phenotype are consistent, but the sub-phenotype CS is not conserved. The results are also confirmed by the observations of various mutational pathways in the resistance evolution of six *ESKAPE* strains(10), revealing a discrete CS phenotype in the pathogens studied. However, CS can be stabilized in different organisms resulting from a common genetic mutational event(12). The convergent kanamycin resistance after resistance evolution in five species of bacteria observed in this study is shaped by high frequency *fusA* mutations. Elongation factor G is a protein with five domains that binds to the 70S ribosome(27) (Figure 2). During the translocation steps, domain IV interacts with the ribosomal A-site, facilitating the movement of mRNA and tRNA from the A site to the P site on the ribosome(28, 29). After kanamycin resistance evolution most *fusA* mutations occurred in domain IV, which may alter the affinity of kanamycin for the ribosomal protein, potentially reducing the drug’s effectiveness.

**Figure 2.**
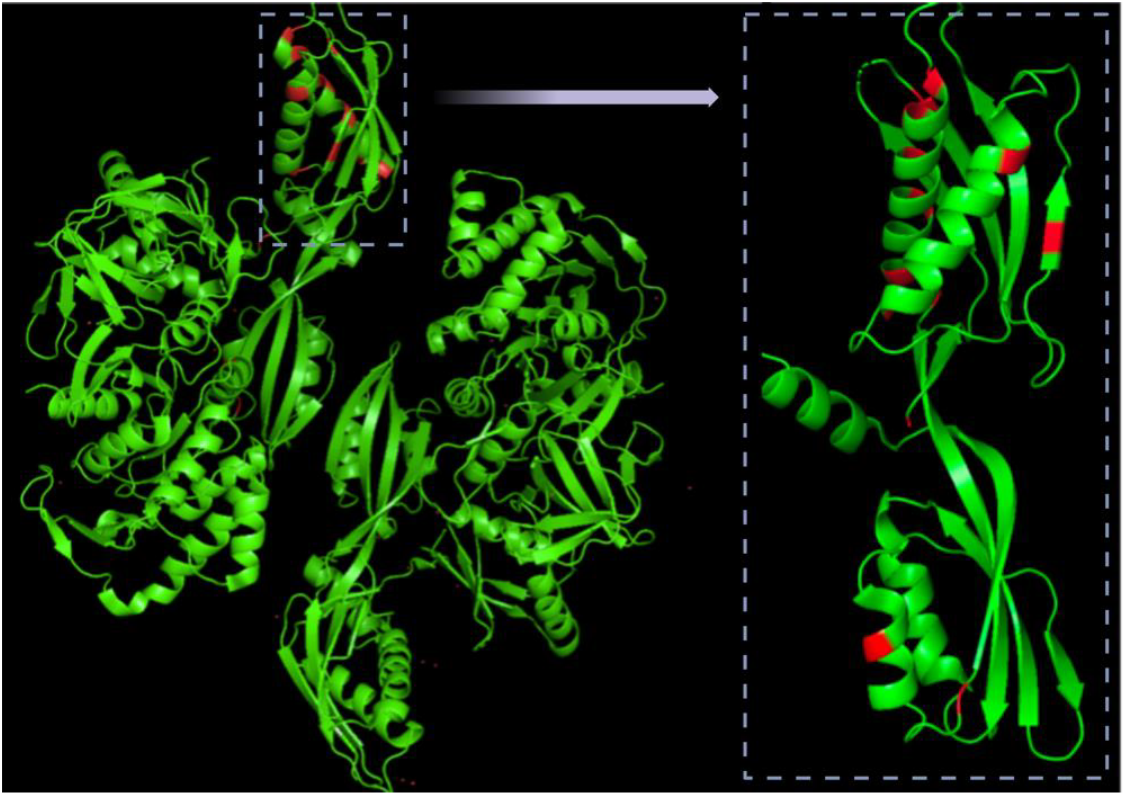
Structural effects of the mutations occurring in the FusA protein of *S. aureus* PDB3ZZT(27).

The mutations that occurred at domain IV in *fusA* result in the same level of resistance to tobramycin and gentamicin in five species of bacteria, indicating that the *fusA* mutations might be the main contributors to aminoglycoside resistance. *FusA* mutations were also acquired in tobramycin evolution in *A. baumannii* and *P. aeruginosa* in biofilm and planktonic condition, implying the mutation are essential for bacteria to survive in different niches(30). As an effect of the common mutations, the strains resistant to chloramphenicol and tetracycline did not induce cross resistance to aminoglycoside antibiotics. Tetracycline and chloramphenicol are bacteriostatic antibiotics and induce less *de novo* resistance than bactericidal compounds(22). This resistance is caused more by cellular adjustments at the expression level than by DNA mutations(23). As a result, they cause more CS, because the adjustments that increase tolerance to bacteriostatic antibiotics apparently increase the sensitivity to bacteriostatic antimicrobials. In addition, the mutations in *fusA* exert a pleiotropic effect(31), which manifested itself in this study as efflux pumps operating in parallel^27^. The *fusA* mutation reduces the reaction to nutritional stress sensors (ppGpp), decreasing heme levels, thereby enhancing oxidative stress and DNA damage. As a result, it increases the susceptibility to β-lactam antibiotics(31), which explains the collateral sensitivity of β-lactam observed in kanamycin resistant strains(30). Based on the CS findings in five resistant strains, we proposed a follow-up treatment line using chloramphenicol/tetracycline, kanamycin and amoxicillin to minimize the effects of induced resistance (Figure. 3).

**Figure 3.**
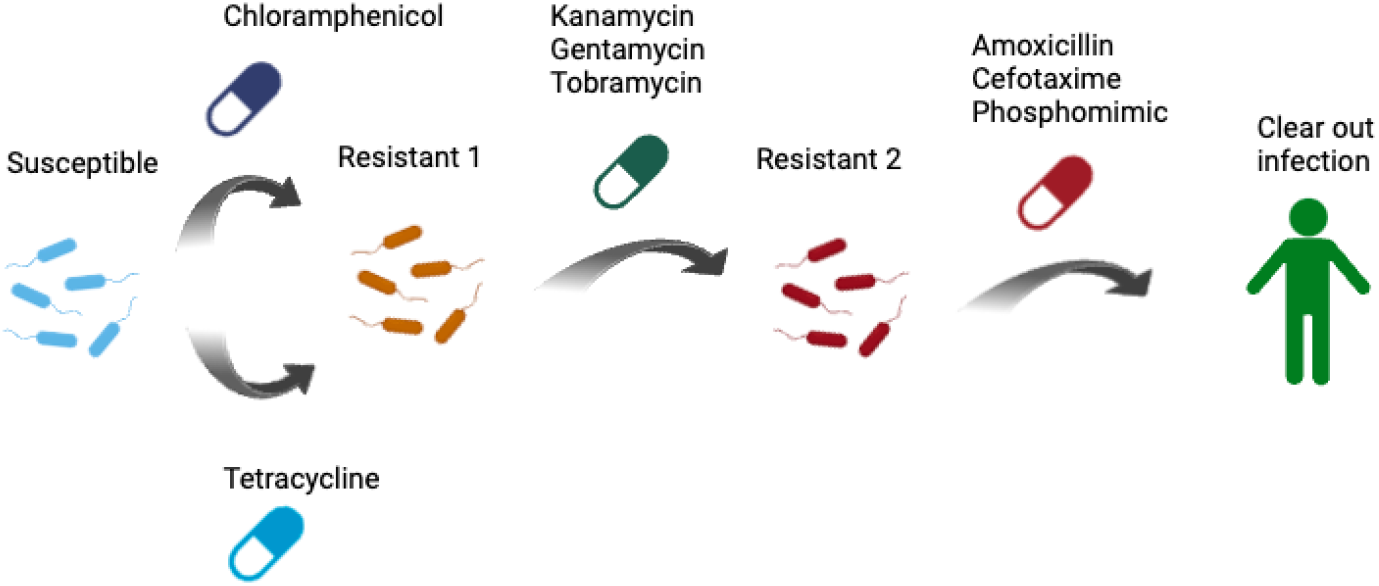
Hypothetical treatment sequence utilizing CS for optimal effectivity.

In conclusion, exposure to bacteriostatic antibiotics increases their sensitivity to aminoglycosides, which in turn make the bacteria more susceptible to beta-lactam antibiotics. Therefore, a theoretical sequence of antimicrobials for persistent infection that develop resistance during treatment can be imagined. Based solely on considerations of resistance, one would arrive at the sequence depicted in figure 3: Start with a bacteriostatic, follow-up with an aminoglycoside and clear the infection with a beta-lactam. Whether this is relevant for the clinic or veterinary practice needs to be investigated.

## Materials and Methods

### Medium, bacterial strains and drugs

The resistant strains were derived from naive wildtypes by exposure to stepwise increasing sublethal concentrations. The wildtype strains were *Salmonella enterica* DSM 9221, *Staphylococcus aureus* DSM 110565, *Enterococcus faecalis* ATCC 4707711, *Acinetobacter pittii* ATCC 33305, *Bacillus subtilis* B168 and *Yersinia enterocolitica* ATCC 9610. The strains acquired resistance against the drugs listed in *Table 1*. The antibiotics were chosen so that each species was exposed to a member of the six classes of antibiotics, but antimicrobials against which inherent resistance existed were avoided. The minimal inhibitory concentration (MIC) measurements(32, 33)and overnight cultures were performed in tryptic soy broth (TSB) medium. Strains were stored at −80°C and streaked onto TSB agar medium for use in overnight cultures. Antibiotics solutions were made from powder stocks (Sigma) and filter sterilized. The antibiotic solutions were kept at −20 °C, or −80 °C for phosphomimic and cefepime, for up to 90 days. Every five days a new batch of amoxicillin was prepared. The antibiotic solutions were stored in the fridge at <4 °C for a maximum of four days.

### De novo resistance acquisition

To develop *de novo* resistance acquisition, the wildtype bacteria were initially exposed to one eights of their MIC for that specific antibiotic. The cell cultures were grown in TSB medium for 24 hours at 200 rpm 37 °C for *B. subtilis, S. aureus, E. faecalis* and *S. enterica*. For *Y. enterocolitica* and *A. pittii*, the strains were incubated at 200 rpm and 30 °C. If the optical density (OD) at an absorbance of 600 nm of the resistant strains was 75% or more compared to the OD of the positive control, the experiment was continued with a two-fold increase of antibiotic concentration with a starting OD=0.1 of the strain. The *de novo* resistance evolution was executed with biological duplicates.

### Minimal inhibitory concentration measurement

To measure the minimal inhibitory concentration (MIC) for the resistant and wildtype strains, the MIC was determined by measuring the absorbance in 96-wells plates (Thermo Fisher Scientific). Overnight cultures were prepared in TSB medium and incubated at 200 rpm 37 °C for *B. subtilis, S. aureus, E. faecalis* and *S. enterica*. For *Y. enterocolitica* and *A. pittii*, the strains were incubated at 200 rpm and 30 °C. The MIC measurements were performed in 96-wells plates with a two-fold dilution of antibiotics per well(32, 33). The complete list of used antibiotics can be found in *Table* 1. The MICs were measured with a minimum of two technical replicates. The bacteria were added per well with an OD of 0.05. Afterwards, the plates were incubated at 200 rpm for 23 hours and 30 °C or 37 °C respectively. Plates were shaken in the plate reader for two minutes and afterwards measured at an absorbance of 600 nanometers.

### Analysis

Heatmaps and graph bars were created using the software GraphPad Prism 9. Collateral sensitivity networks were mapped using the software Cystoscope 3.10.0. To calculate the degree of collateral sensitivity or cross-resistance, the following formula was used:

log2 *MIC*(*resistant strain*) divided by (*MIC*(*wildtype strain*)).

A negative outcome represents collateral sensitivity and a positive outcome cross-resistance compared to their wildtype.

### Mutation analysis

The *fusA* mutations were analyzed in de novo resistant strains after whole genome sequencing(1). All of *fusA* mutations found in the six species used were aligned to the FusA protein of *S. aureus* PDB3ZZT(27).

## Acknowledgements

We thank Selina van Leeuwen for her assistance in DNA sequencing. We appreciate the efforts made by students Nimo Annor, Gioia Schaar in this project. Xinyu Wang acknowledges the China Scholarship Council supporting his PhD scholarship.

